# Establishment of well-differentiated camelid airway cultures to study Middle East respiratory syndrome coronavirus

**DOI:** 10.1101/2021.11.10.468038

**Authors:** Mitra Gultom, Annika Kratzel, Jasmine Portmann, Hanspeter Stalder, Astrid Chanfon Bätzner, Hans Gantenbein, Corinne Gurtner, Nadine Ebert, Horst Posthaus, Patrik Zanolari, Stephanie Pfaender, Volker Thiel, Ronald Dijkman

## Abstract

In 2012, Middle East respiratory syndrome coronavirus (MERS-CoV) emerged in Saudi Arabia and was mostly associated with severe respiratory illness in humans. Dromedary camels are the zoonotic reservoir for MERS-CoV. To investigate the biology of MERS-CoV in camelids, we developed a well-differentiated airway epithelial cell (AEC) culture model for *Llama glama* and *Camelus bactrianus*. Histological characterization revealed progressive epithelial cellular differentiation with well-resemblance to autologous *ex vivo* tissues. We demonstrate that MERS-CoV displays a divergent cell tropism and replication kinetics profile in both AEC models. Furthermore, we observed that in the camelid AEC models MERS-CoV replication can be inhibited by both type I and III interferons (IFNs). In conclusion, we successfully established camelid AEC cultures that recapitulate the *in vivo* airway epithelium and reflect MERS-CoV infection *in vivo*. In combination with human AEC cultures, this system allows detailed characterization of the molecular basis of MERS-CoV cross-species transmission in respiratory epithelium.

## Introduction

Middle East respiratory syndrome coronaviruses (MERS-CoV) emerged in 2012 as the causative agent of severe viral pneumonia in humans. To date, more than 2500 laboratory confirmed cases have been reported, with a case fatality rate of 34% (1). MERS-CoV is a zoonotic pathogen that is intermittently transmitted from dromedary camels to humans leading to local outbreaks with limited human-to-human transmission. Sero-surveillance indicates that MERS-CoV is enzootic in dromedary camels in the Arabian Peninsula and Africa, and in contrast to humans, only causes a mild respiratory tract infection in camelids (2–4). Interestingly, the majority of zoonotic infections in humans, as well as higher sero-prevalence of camel exposed workers, are more frequently observed in the Middle East compared to Africa (5,6). Possible reasons for this pattern might lay in the genetic differences of the virus across the two regions, as well as cultural discrepancies or a lower awareness and surveillance in Africa (5,7–9). Nevertheless, travel-related cases in 27 countries, as well as nosocomial outbreaks in hospitals, such as in South Korea in 2015 (186 cases and 36 deaths), have been reported and highlight the need for continuous surveillance to mitigate future epidemics (10).

The anatomical distance of the conducting airways is markedly different between camelids and humans (11). Concordantly, viral shedding during MERS-CoV infection in humans and camelids is dissimilar, as exemplified by the detection of relatively high levels of MERS-CoV in the upper respiratory tracts of infected camelids, as opposed to humans, in which the infection is restricted to the lower respiratory tract (12–14). This can be partly explained by the different distribution of the functional receptor – serine exopeptidase Dipeptidyl Peptidase-4 (DPP4) – for MERS-CoV in humans and camelids, which likely influences the limited human-to-human transmission (12,15). However, the observed discrepancy in the clinical resolution between MERS-CoV infected humans and camelids also suggests that other host determinants, such as the innate immune system, might be of importance.

To facilitate investigations that are focused on the molecular basis underlying the pathogenesis discrepancy of MERS-CoV in humans and camelids, we develop a pseudostratified airway epithelial cell (AEC) culture model for llama (*Llama glama*) and Bactrian camel (*Camelus bactrianus*), analogous to the human AEC culture model. Histological and functional characterization revealed that both camelid AEC culture models closely resemble the *in vivo* morphology, actively respond to IFNs, and are permissive to MERS-CoV. These three characteristics illustrate that the established camelid AEC model allows detailed future comparative studies on virus-host interactions in human and camelid respiratory epithelium.

## Results

### Establishment of camelid airway epithelial cell cultures

To generate *in vitro* models that potentially serve as surrogates to characterize MERS-CoV – host interaction at the main replication site in the host reservoir, we sought to establish well-differentiated airway epithelial cell cultures from camelids, analogous to human AEC cultures. Unfortunately, there was no tracheobronchial tissue available from *Camelus Dromedarius* during the entire study period (2014 – 2021), due to national import and export restrictions. As both Bactrian camels and llama are also susceptible to MERS-CoV infection, we chose to isolate primary epithelial cells from post-mortem tissue from tracheobronchial regions of *Camelus bactrianus* (1 donor) and *Lama glama* (2 donors) and propagated them using a pre-established protocol (16). Following the isolation and expansion, epithelial cells from the old and new world camelids were seeded on semi-permeable cell culture inserts. Once cells reached confluency, the cultures were air-lifted to establish an Air-liquid interface (ALI) to allow for cellular differentiation. During the differentiation process, the development of the camelid AEC cultures was monitored by immunofluorescence analysis with 7-day intervals for a total duration of 28 days.

This revealed a progressive ciliary development in both Bactrian camel and llama AEC cultures that reached a plateau after three weeks for the llama AEC cultures. For the Bactrian camel AEC cultures, the overall number of ciliated cells was slightly lower (**Figure 1A-C**). Nonetheless, for both species tight junction formation seemed to stabilize two weeks after ALI establishment as indicated by the trans-epithelial electrical resistance (TEER) measurement and condensed hexagonal architecture of the tight junction marker Zona Occludens 1 (ZO-1) (**Figure 1A, D**). In addition to the quantitative measurements, camelid AEC cultures were histologically compared with autologous *ex vivo* tissue from the corresponding anatomical region. These vertical histologic sections demonstrated that after 28 days of differentiation the camelid AEC cultures formed pseudostratified layer of epithelial cells (**Figure 1E**). Combined these results demonstrate that the well-differentiated camelid AEC cultures exhibit morphological properties resembling the Bactrian camel and llama tracheobronchial respiratory epithelium.

**Figure 1.**
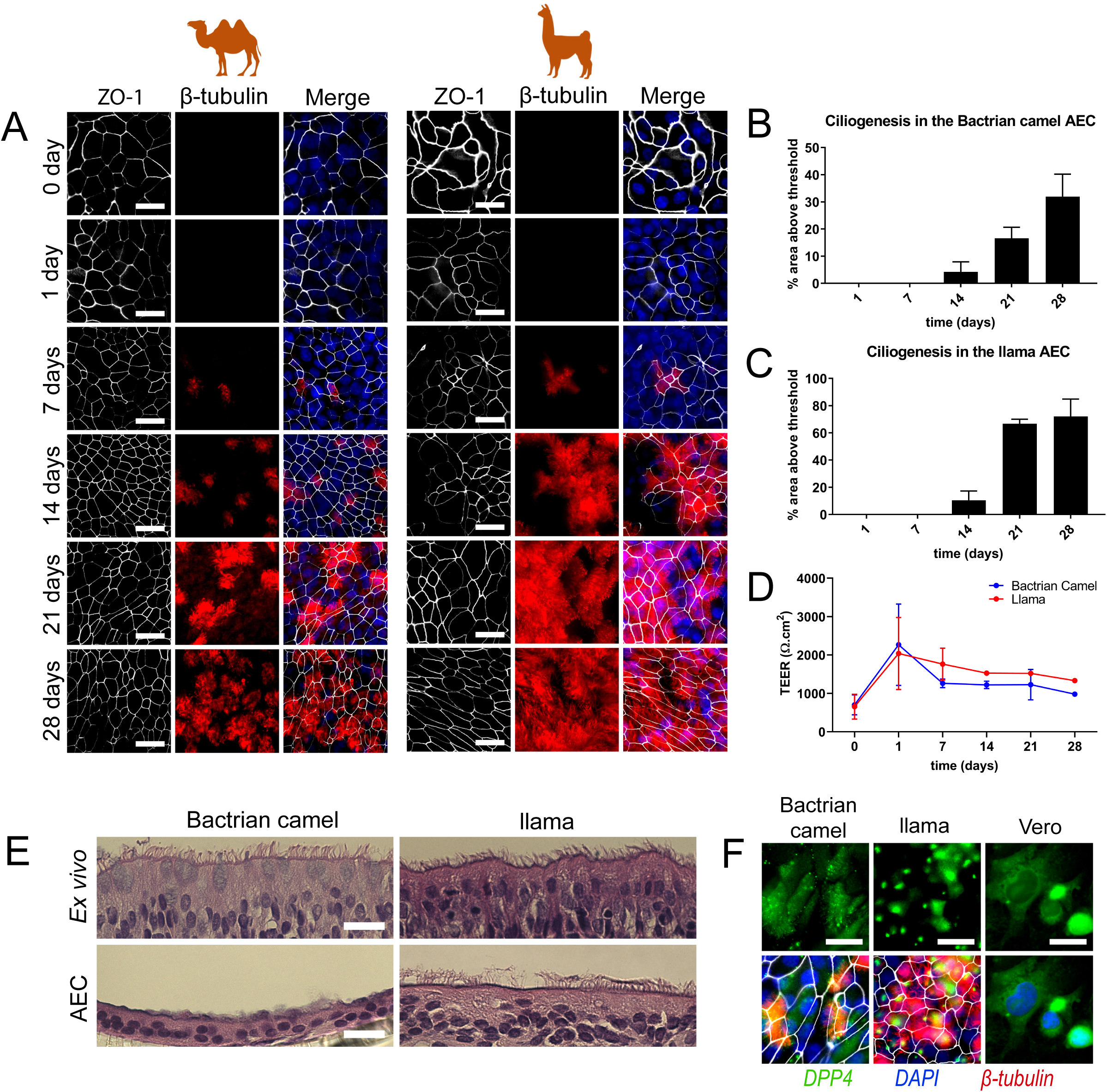
Establishment and characterization of Bactrian camel and llama AEC cultures. (**A**) Immunofluorescence analysis showing the development of tight-junctions (ZO-1, white) and ciliogenesis (β-tubulin, red) in Bactrian camel and llama AEC cultures over time from 1-day to 4 weeks post ALI exposure. The cells were counterstained with DAPI (blue) to visualize the nuclei. (**B**) and (**C**) Ciliogenesis quantification of camel and llama AEC cultures overtime, respectively. Ciliation was quantified by measuring the area above a fluorescence intensity threshold of five random images acquired per condition. (**D**) Transepithelial electrical resistance (TEER) measurement of camel and llama AEC cultures overtime during the differentiation. (**E**) Epithelial morphology of *ex vivo* tissues (upper panel) and well-differentiated camel and llama AEC cultures (lower panel). (**F**) DPP4 expressions of well differentiated camel and llama AEC cultures, with Vero cells as a positive control. Scale bar is 20 μm.

### Efficient MERS-CoV replication in Camelid AEC cultures

Following the establishment of the camelid AEC cultures, we assessed the expression and distribution of the functional receptor for MERS-CoV in formalin-fixed AEC cultures using a polyclonal antibody against DPP4. As a positive control, we included the Vero E6 cell line, which is known to express DPP4 (15,17). This revealed that DPP4 could readily be detected in both Bactrian camel and llama AEC cultures (**Figure 1F**). Of note, the Bactrian camel DPP4 was predominantly distributed at the apical surface of non-ciliated cell populations, while in llama AEC cultures, DPP4 expression is mainly restricted to the apical surface of the ciliated cell population (**Figure 1F**). To determine whether both camelid AEC models are susceptible to MERS-CoV, we inoculated well-differentiated Bactrian camel and llama AEC cultures with 4000 PFU of MERS-CoV (MERS-CoV EMC/2012) at 37 °C. At 2 hours post-infection (hpi), the apical surface was washed three times with HBSS. Subsequently, virus progeny release was monitored every 24h for the duration of 96 hours by virus titration and qRT-PCR. Interestingly, in both Bactrian camel and llama AEC cultures we observed efficient, albeit dissimilar, MERS-CoV replication profiles. In Bactrian camel AEC cultures MERS-CoV readily reached a plateau at 24 hpi after which the amount of infectious progeny virus declined overtime (**Figure 2A, C**). In contrast, the overall MERS-CoV replication kinetics in llama AEC cultures was delayed, as MERS-CoV reached the highest titer at 96 hpi (**Figure 2B, D**).

**Figure 2.**
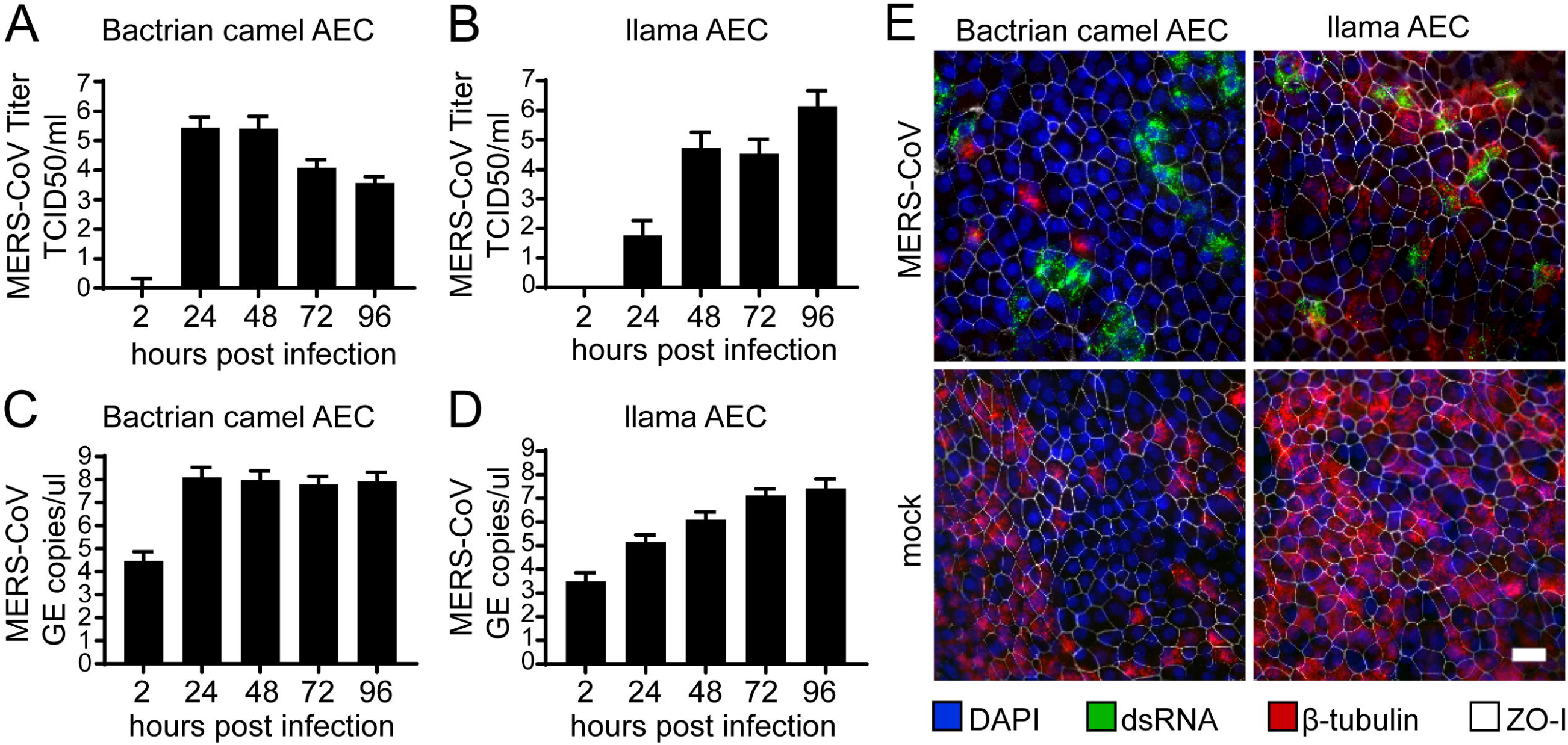
Efficient MERS-CoV replication in camelid AEC cultures. MERS-CoV titer in TCID50/ml released from camel (**A**) and llama (**B**) AEC cultures’ apical side from 24 to 96 hours post infection depicted in a log10 scale. Apical virus release in Bactrian camel (**C**) and llama (**D**) AEC cultures measured by quantitative reverse transcription PCR. (**A, C**) Results from three independent replications from one biological donor are shown. (**B, D**) Results from six independent replications from two biological donors are displayed. Graphs are shown in mean and standard deviation. (**E**) Immunofluorescence staining of MERS-CoV-infected camel and llama AEC cultures 48 post infection are depicted. Double-stranded RNA is shown in green, β-tubulin in red, ZO-1 in white and DAPI in blue. Scale bar is 20 μm.

Following the replication kinetics, we also assessed the viral cell tropism with immunofluorescence analysis at 48 hpi using an antibody against double stranded RNA (dsRNA), as a surrogate marker for active viral replication. This highlighted that in both Bactrian camel AEC cultures dsRNA was mainly observed in non-ciliated cell populations, similar as in human AEC cultures (**Figure 2E, left panels**) (18). In contrast, in the llama AEC cultures dsRNA-positive cells were predominantly co-stained with the cellular ß-tubulin marker used to detect ciliated cells (**Figure 2E, right panels**). Altogether, these results demonstrate that both camelid AEC cultures support efficient MERS-CoV replication and indicate that the viral cell tropism coincides with the DPP4 distribution in the AEC models from both camelid species.

### Recombinant IFN inhibits MERS-CoV replication

We have previously demonstrated that MERS-CoV replication can be reduced upon exogenous type I and III IFN treatment in human AEC cultures (18). However, prior to assessing whether MERS-CoV replication can be reduced upon exogenous type I and III IFN treatment in both camelid AEC cultures we first determined whether the cell-intrinsic innate immune system is functional. For this, we stimulated camelid AEC cultures with recombinant pan-species type I IFN, human type III IFN, and synthetic dsRNA (poly-I:C) and analyzed the induction of several host transcripts at 6 h and 12 h post-treatment. This revealed that stimulations by exogenous type I and type III IFNs lead to the induction of canonical ISGs, such as Mx1, CXCL10, and RIG-I in both camelid AEC cultures (**Figure 3A, B**). Stimulation with poly I:C, a surrogate for active virus replication, resulted in the induction of both ISGs and chemokines (**Figure 3C**), signifying that both the sensing and signaling arms of the cell-intrinsic innate immune system seem to be intact in both camelid AEC models.

**Figure 3.**
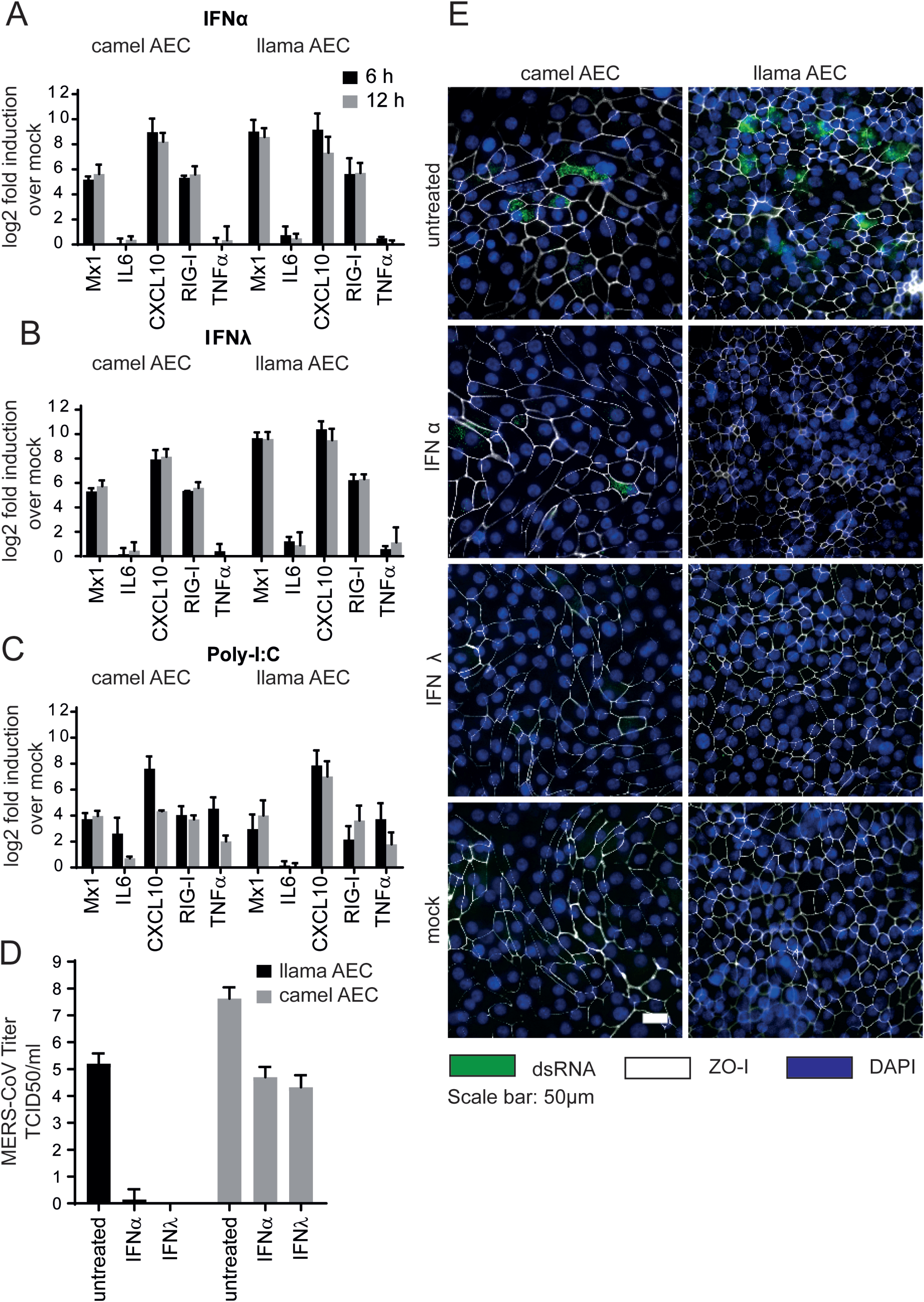
IFN treatment efficiently induces innate immune response and reduces MERS-CoV replication in camelid AEC cultures. MX1, Interleukin-6 (IL6), CXCL10, RIG-I and TNF-α expression in log_2_ fold induction over mock in camel and llama AEC cultures is displayed 6- and 12- hours post stimulation with type I IFN (**A**), type III IFN (**B**), and poly I:C (**C**) treatment. (**D**) MERS-CoV titers in TCID50/ml released from camelid AEC cultures’ apical side 48 hours post infection is shown in presence and absence of type I and III IFN pretreatment. (**E**) Immunofluorescence staining of type I and III IFN pretreated camelid AEC cultures is shown 48 hours post infection. Double-stranded RNA is shown in green, ZO-1 in white and DAPI in blue. Scale bar is 50 μm.

After having established the functionality of the innate immune system in the camelid AEC cultures, we assessed whether exogenous type I and III IFN treatment could reduce MERS-CoV replication in the camelid AEC cultures, similar as in human AEC cultures. For this, camelid AEC cultures were pre-stimulated with exogenous type I or III IFNs for 18 hours prior to MERS-CoV infection. Forty-eight hours post-infection, we observed that in the Bactrian camel AEC cultures apical progeny virus release was reduced by 3.5 to 4 logs by type I and type III IFNs, respectively. Interestingly, for the llama AEC cultures we observed that both type of IFNs markedly reduced MERS-CoV replication (**Figure 3D**). These results were corroborated by immunofluorescence analysis at 48h, in which the number of dsRNA positive cells are reduced in IFN treated cultures compared to untreated cultures (**Figure 3E**). This indicates that, like human AEC cultures, MERS-CoV replication can be reduced in both camelid AEC cultures upon type I and III IFN pretreatments. Combined, these results demonstrate that the established camelid AEC models facilitates future experimental work to dissect fundamental virus – host interactions within the camelid respiratory epithelium.

## Discussion

In the current study, we have successfully established well-differentiated AEC cultures from Bactrian camel and llama as surrogate *in vitro* models to study MERS-CoV in the camelid hosts. We show that the camelid AEC cultures are well-differentiated, possess a functional innate immunity, and support efficient MERS-CoV replication. We observe the cell tropisms and replication kinetics profiles during MERS-CoV infection are dissimilar among the old-world and new-world camelid AEC cultures. In addition, we demonstrated that pretreatments with exogenous type I and III IFNs were able to reduce MERS-CoV viral replication in both camelid AEC cultures. Our data suggest that well-differentiated camelid AEC cultures can serve as a more biologically relevant system that morphologically and functionally resembles the natural site of infection of MERS-CoV in camelids. This approach can help to circumvent the needs of certain animal experiments, especially since experiments using large animals such as camelids are difficult to conduct due to their limited availability and logistical requirements.

While dromedary camels are shown as the main zoonotic reservoir for MERS-CoV, susceptibility of both Bactrian camel and llama to MERS-CoV by either natural or experimental infection has been previously reported (14,19–21). Although both species are genetically closely related, we show that the replication profile as well as cell tropisms of MERS-CoV are different in Bactrian camel and llama, in line with those observed results in experimentally infected camels and llamas (13,14,19). Interestingly, as the natural habitat of Bactrian camels shows a higher overlap to that of dromedary camels, the reservoir for MERS-CoV compared to llamas, MERS-CoV might evolutionarily favor and be better adapted to the Bactrian camel (21). Differences in MERS-CoV cell tropism can be explained by the distinct MERS-CoV receptor distribution between camel and llama AEC cultures. Nevertheless, differential DPP4 distribution along the respiratory tract of camelids and other ungulates which correlates to their susceptibility has also been described (12,19,22). However, it was previously shown that the presence of DPP4 alone does not always translate to susceptibility in other animals *in vivo*, highlighting the importance of investigating the role of other host determinants in the outcome of MERS-CoV infection (19,23).

In this study, we demonstrate that camelid AEC cultures are responsive to type I and III IFN stimuli and that pretreatment with exogenous IFNs can reduce MERS-CoV replication, similar to previously observed results in primary human AEC cultures (18). However, despite the close evolutionary relationship between llamas and Bactrian camels, we did observe a species-specific difference in the efficacy of inhibiting MERS-CoV replication upon exogenous stimulation with type I and III IFNs. This suggests potential differences in IFN receptor distribution and/or downstream signaling cascades inducing the expression of ISGs tempering MERS-CoV replication. These results, together with the previously reported *in vivo* data of MERS-CoV infection in alpacas, suggest dominant role for type I and III IFNs in camelids (24). Moreover, since both recombinant type I and III IFNs also efficiently inhibited MERS-CoV in human AEC cultures it would be worth to evaluate the therapeutic potential of IFNs towards MERS-CoV, as well as further investigating MERS-CoV interaction with IFN-related pathways in different host species (17). Such analyses can now be performed using the camelid AEC cultures in conjunction with the analogous human AEC cultures, to provide detailed information on crucial virus-host innate immune response dynamics in both the natural and zoonotic hosts.

In summary, our results demonstrate that these cultures can serve as a biologically relevant model to characterize fundamental molecular virus-host interactions of MERS-CoV at the natural site of infection in camelids. Altogether, the established camelid AEC culture system, in combination with human AEC cultures, facilitates future detailed characterization of the molecular basis of the pathogenesis discrepancy of MERS-CoV in humans and camelids in a biologically relevant system.

## Materials and Methods

### Establishment of camelid AEC cultures

Tracheobronchial epithelial cells from Bactrian camel and llama were isolated from post-mortem tracheobronchial tissue, obtained in collaboration with the veterinary hospital of the University of Bern that euthanized their animals for diagnostic purposes. Isolation and culturing were performed as previously described (16). Modifications to the composition of the ALI medium were introduced, in which the concentration of the EGF was increased to 5 ng/ml. Both camel and llama ALI cultures were maintained at 37 °C in a humidified incubator with 5% CO_2_. During the development of differentiated camelid ALI cultures (3-4 weeks), media was changed every 2-3 days. During the ALI differentiation stage, inserts were fixed at 7-day intervals from the day of ALI exposure to 4-week post-ALI to monitor the development. TEER resistance was measured every seven days.

Histological examination of both *ex vivo* tissues and well-differentiated camelid AEC cultures were done by formalin fixation and staining with haematoxylin and eosin (HE) according to standard histological techniques. The sections were observed and visualized using an EVOS FL Auto 2 imaging system (Thermo Fisher Scientific). Acquired images were processed with Fiji software package v1.53 (25).

### Conventional cell lines

Human hepatoma (Huh7) cell line (kindly provided by Volker Lohmann) was propagated in Dulbecco’s Modified Eagle Medium (DMEM), supplemented with 10% heat-inactivated fetal bovine serum, 1% nonessential amino acids, 100 μg/mL of streptomycin, 100 IU/mL of penicillin, and 15 mM of HEPES. Cells were maintained at 37 °C in a humidified incubator with 5% CO_2_. Huh7 cell line was confirmed to be of human origin without contamination, matching the reference DNA of the cell line Huh7 (Microsynth reference, Mic_ 152021) with 96.7% and the DNA profile of Huh7 (Cellosaurus, RRID:CVCL_0336) with 90%.

### Virus Infection

Well-differentiated camelid AEC cultures were infected with 4000 PFU of MERS-CoV (strain EMC/2012, propagated on Huh7 cells diluted in Hanks balanced salt solution (HBSS, Gibco). The cells were washed with 100 μl of HBSS prior to infection (18). The virus was inoculated via the apical side. Virus-infected and control AEC cultures were incubated at 37 °C in a humidified incubator with 5% CO_2_. After the inoculation, inoculum was removed, and the apical surfaces were rinsed three times with HBSS, where the third washes were collected as 2 h timepoint. Progeny virus release was monitored with 24-hour intervals for a total duration of 96 hours, through the application of 100 μl of HBSS onto the apical surface, incubated 10 min prior to the collection time point. The collected apical washes were diluted 1:1 with virus transport medium (VTM) and stored at −80°C for later analysis. Following the collection of the apical washes, the basolateral medium was exchanged with fresh ALI medium. Each experiment was repeated as three independent biological replicates. For camel, biological replicates were generated from one donor that whereas for llama, each replicate represents a different biological donor.

### Interferon and poly-I:C stimulation

To analyze the response of camelid AEC cultures to IFN stimulations, both Bactrian camel and llama AEC cultures were treated with recombinant universal type I and III IFNs (100 IU/ml; Sigma Aldrich) or recombinant type III IFN (100 ng/ml) for 6 and 12h at 37 °C from the basolateral side (26). For poly-I:C stimulation, AEC cultures were treated with 10 μg poly-I:C (Sigma Aldrich) in 50 μl of HBSS from the apical sides for 6 and 12h at 37 °C. Thereafter, total cellular RNA from the pretreated cells were isolated with the NucleoMag RNA kit (Macherey-Nagel) according to the manufacturer’s guidelines on a Kingfisher Flex Purification system (Thermo Fisher Scientific). The quantity of the RNA was determined using NanoDrop (Thermo Fisher Scientific). From the total RNA, cDNA was synthesized using MMLV reverse transcriptase kit (Promega) according to the manufacturers’ protocol with random primers (Promega). Two microliters of diluted cDNA were amplified with SYBR™ Green PCR Master Mix (Thermo Fisher Scientific) according to the manufacturer’s protocol, using primers targeting five different interferon-stimulated genes (ISGs) transcripts (**Table. 1**). GAPDH was used as the reference gene. Measurements and analysis were performed with the Applied Biosystems™ 7500 Fast Dx Real-Time PCR Systems and associated software (Applied Biosystems). Relative gene expression was calculated using the 2^ΔΔCt^ method (27). Data are shown as fold induction of IFN-treated samples compared to those of untreated controls.

**Tabel 1.**
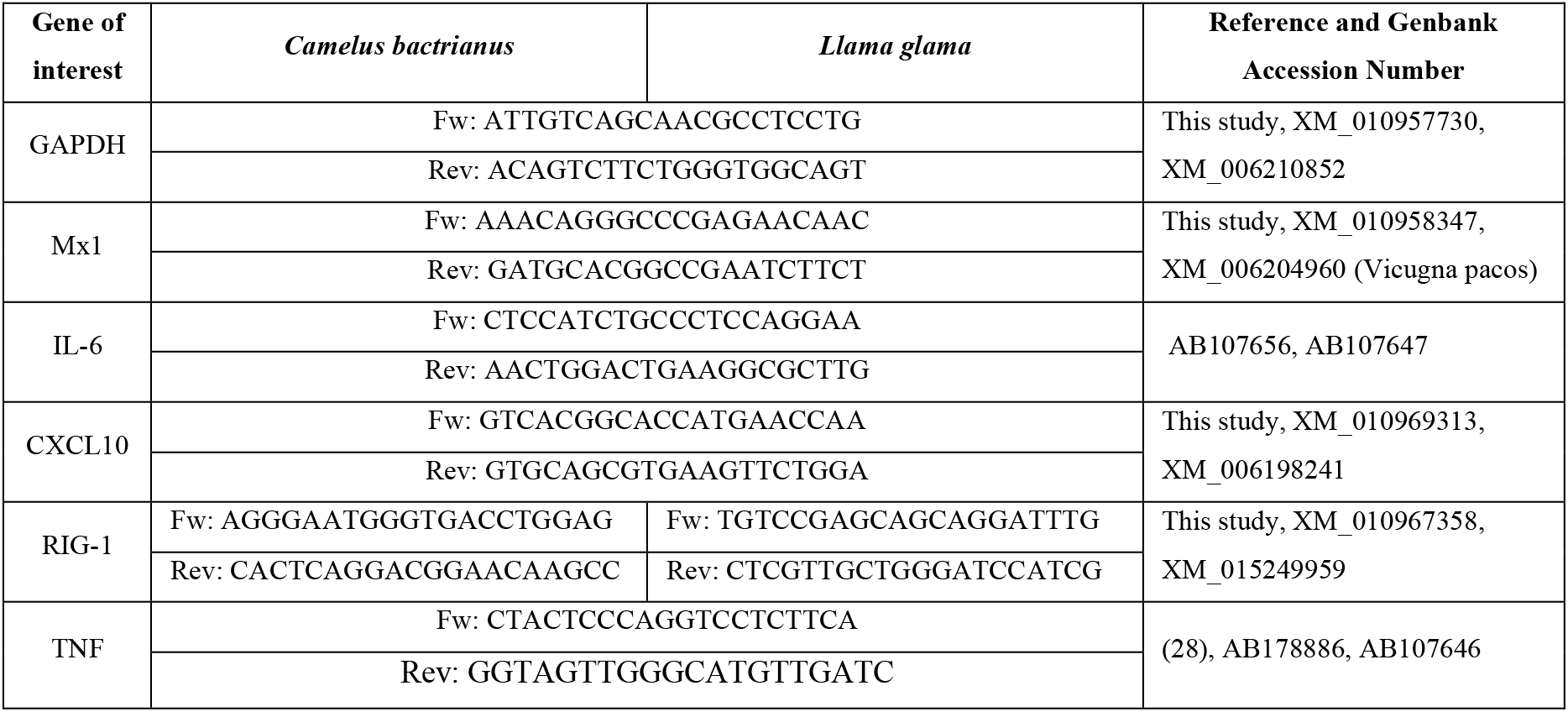
List of primers used to identify the ISGs in Bactrian camel and llama AEC cultures.

To examine the influence of IFN pretreatment on MERS-CoV infection in camelid AEC cultures, the cells were pretreated with type I and III IFNs (100 IU/ml and 100 ng/ml, respectively) 18h prior to MERS-CoV infection at 37 °C. The basolateral medium containing type I or type III IFN was removed and replaced with medium without exogenous IFN before infection. Untreated Bactrian camel and llama AEC cultures were used as controls. Progeny virus released on the apical sides was collected with 100 μl of HBSS at 48 hpi.

### Virus titration

The 50% tissue culture infectious dose (TCID50) per milliliter of supernatant was determined by inoculating Huh7 cells with serially diluted apical washes at indicated hours post-infection. 72 h post-inoculation, cytopathic effect (CPE) was visualized using crystal violet, and TCID50 per milliliter was calculated by the Spearman-Kärber algorithm 72h as previously described (29).

### qRT-PCR of MERS-CoV

qRT-PCR method was used to determine virus replication. Viral RNA was isolated from the supernatant at indicated hours post infection using the NucleoMag Vet Kit (Macherey Nagel) and a Kingfisher Flex Purification System (Thermo Fisher Scientific) according to manufacturer’s guidelines. Extracted RNA was amplified using TaqMan™ Fast Virus 1-Step Master Mix (Thermo Fisher Scientific) according to the manufacturers’ protocol. Primers used for detection of MERS-CoV targeting regions upstream of the E gene (upE) are listed on **Table 2** (Genbank accession numbers NC038294 and MG923481) (30). A serial dilution of *in vitro* transcribed MERS-CoV RNA (kindly provided by Victor Corman) was used as a reference (30). Measurements and analysis were performed with the Applied Biosystems™ 7500 Fast Dx Real-Time PCR Systems and associated software (Applied Biosystems).

**Table 2.**
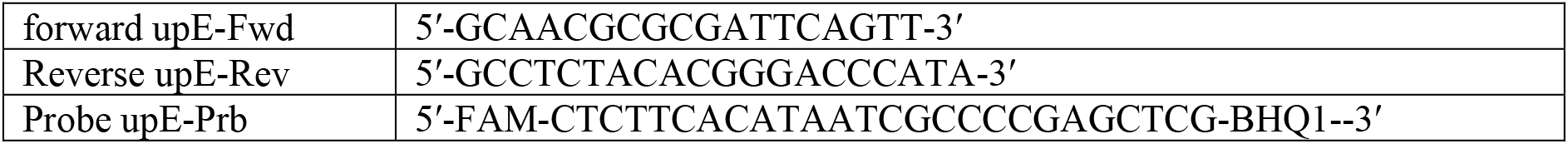
Primers used to analyze MERS-CoV replication in Bactrian camel and llama AEC cultures.

### Immunofluorescence analysis

Cells were fixated with 4% formalin for immunofluorescence analysis. Fixated cells were permeabilized in PBS supplemented with 50 mM NH_4_Cl, 0.1% (w/v) Saponin, and 2% (w/v) Bovine Serum Albumin and stained with a mouse monoclonal antibody against dsRNA (SCICONS, clone J2). Alexa-Fluor 488-labeled donkey-anti mouse IgG (H+L) (Jackson Immunoresearch, 715-545-150) was used as a secondary antibody. To visualize the cellular receptor DPP4, a rabbit polyclonal antibody against DPP4 (Abcam, ab28340) was used. Subsequently, Alexa-Fluor 488-labeled donkey-anti rabbit IgG (H+L) (Jackson Immunoresearch, 711-545-152) was used as a secondary antibody. Alexa-Fluor® 647-labelled rabbit anti β-tubulin IV (Cell Signalling Technology, 9F3) and Alexa-Fluor® 594-labelled mouse antibody against ZO-1 (Thermo Fisher Scientific, 1A12) were used to visualize cilia and tight junctions, respectively. Cells were counterstained using 4’ ,6-diamidino-2-phenylindole (DAPI, Thermo Fisher Scientific) to visualize the nuclei. Images were acquired using an EVOS FL Auto 2 Imaging System, using 40x air objectives. Brightness and contrast were adjusted identically to the corresponding controls using the Fiji software packages and figures were assembled using FigureJ (25,31). Quantification of the ciliation was done on five randomized fields of view of β-tubulin-stained inserts acquired at 40x objective by measuring cilia-positive area above the threshold.

## Acknowledgements

This work was funded by research grants from the European Commission (Marie Sklodowska-Curie Innovative Training Network “HONOURS”; grant agreement No 721367), the Federal Ministry of Education and Research (BMBF; grant RAPID, #01KI1723A), and the Swiss National Science Foundation (SNSF grants 31CA30_196062).

